# A Complete Set of Equations for a Computational Model of Electrolyte and Water Transport along the Nephrons in a Mammalian Kidney

**DOI:** 10.1101/2022.09.23.509286

**Authors:** Anita T. Layton

## Abstract

To investigate kidney under different physiological and pharmacological conditions, we have developed and applied computational models of electrolyte and water transport along nephrons in a kidney. The models predict spredict luminal fluid flow, hydrostatic pressure, membrane potential, luminal and cytosolic solute concentrations, and transcellular and paracellular fluxes through transporters and channels. The complete set of model equations are presented here.

## 1 Introduction

We present a computational model of electrolyte and water transport along nephrons in a mammlian kidney. Typically, superficial nephrons are one of the two main classes of nephrons in a rodent kidney, the other being the juxtamedullary nephrons. The superficial nephrons turn at the boundary of the outer and inner medullas, whereas the juxtamedullary nephrons enter the renal medulla. In a rat kidney, for example, the superficial and juxtamedullary nephrons account for approximately 1/3 and 2/3 of the total nephron population, respectively. In a mouse kidney, the ratio of superficial and juxtamedullary nephrons is 82:18.

The model nephron consists of a number of segments: the proximal tubule, loop of Henle, distal convoluted tubule, connecting tubule, and collecting duct. The water and solute transport processes along each nephron segment are simulated by a number of epithelial cell models connected in series. Each cell model includes the apical (lumen-facing) and basolateral (interstitium-facing) membranes, and is separated from the adjacent cell by a paracellular space. A schematic diagram of the model nephron is shown in Fig. 1.

**Figure 1:**
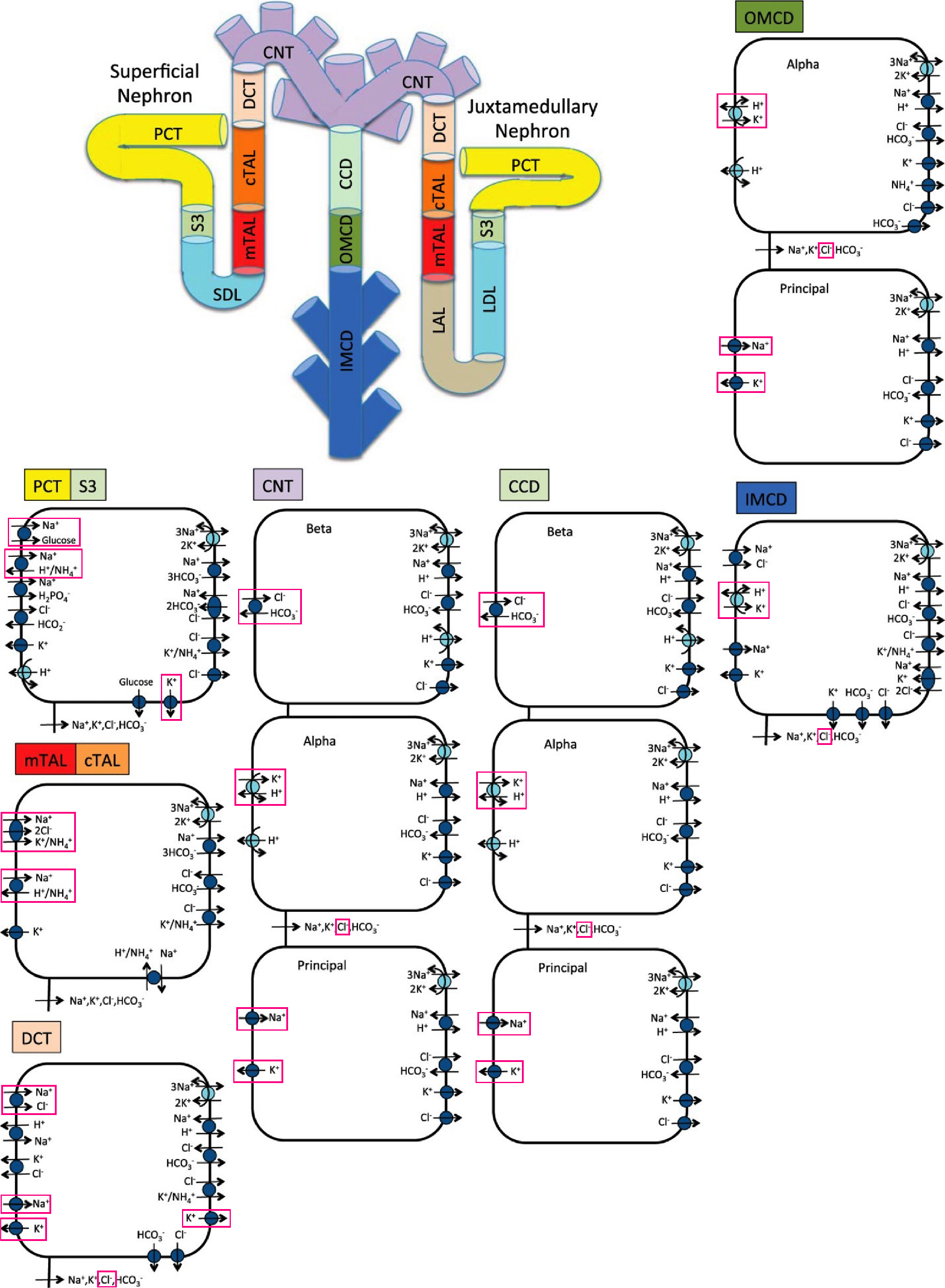
Schematic diagram of the nephron system (not to scale). The model includes one representative superficial nephron and five representative juxtamedullary nephrons, each scaled by the appropriate population ratio. Only the superficial nephron and one juxtamedullary nephron are shown. Along each nephron, the model accounts for the transport of water and 15 solutes (see text). The diagram displays only the main Na^+^, K^+^, and Cl^−^ transporters. Transporters and channels that are assumed regulated by the circadian clock are highlighted in red. PCT, proximal convoluted tubule; SDL, short or outer medullary descending limb; mTAL, medullary thick ascending limb; cTAL, cortical thick ascending limb; DCT, distal convoluted tubule; CNT, connecting tubule; CCD, cortical collecting duct; OMCD, outer-medullary collecting duct; IMCD, inner medullary collecting duct; LDL, thin descending limb; LAL, thin ascending limb.

The model simulates the transport of water and 15 solutes: 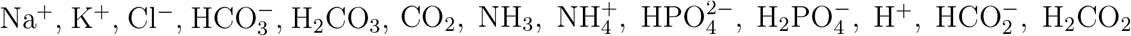, urea, and glucose. The model is formulated for steady state and predicts luminal fluid flow, hydrostatic pressure, luminal fluid solute concentrations, cytosolic solute concentrations, and membrane potential. The model also predicts transcellular and paracellular fluxes by determining water and solutes that move between two adjacent compartments. Model parameters under baseline conditions can be found in Ref. [12] and the references therein. The present study allows for rhythmic variations in some of the hemodynamics and transport parameters.

## 2 Conservation equations for the cellular and paracellular compartments

The cellular and paracellular (i.e., lateral) compartments are assumed to be non-compliant. Thus, water conservation in the cellular and paracellular compartments (denoted by subscripts ‘C’ and ‘P’, respectively) of tubular segment *i* is given by

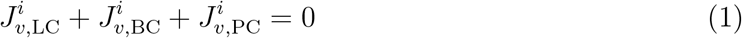

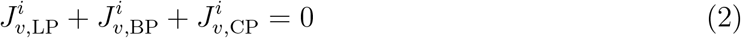

where the subscripts ‘L’ and ‘B’ denote lumen and blood (i.e., interstitium), respectively. Schematic cell models corresponding to individual nephron segments are shown in Fig. 1. The cellular and paracellular compartments (‘C’ and ‘P’) correspond to the cellular interior and the paracellular space shown underneath each cell model. The luminal and blood compartments (‘L’ and ‘B’) are the space on the left and right, respectively, of each cell model. In the above notations, water flux 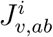 is taken positive from compartment *a* to *b*.

Conservation of a non-reacting solute *k* is given by

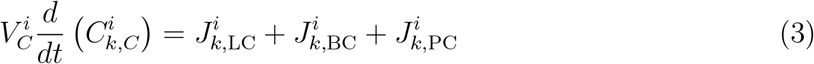

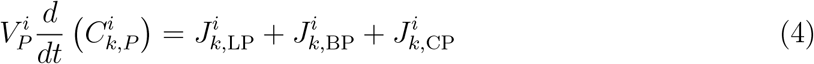

where 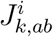 denotes the transmembrane flux of solute *k* from compartment *a* to *b*. 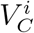 and 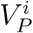 denote the volume of the cellular (‘*C*’) and paracellular (‘*P* ‘) compartments, respectively.

One of the kidney’s functions is to maintain acid-base balance, in part by excreting hydrogen ions into the urine and reabsorbing bicarbonate from the urine. For the reacting solutes, conservation is applied to the total buffers:

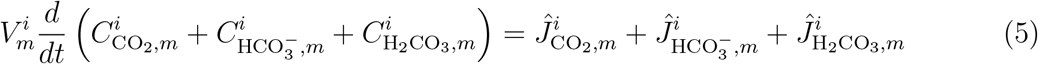

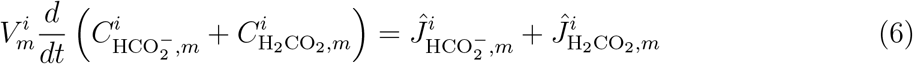

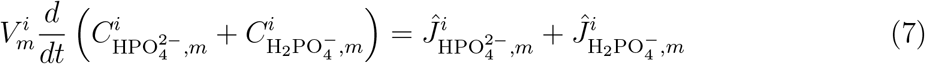

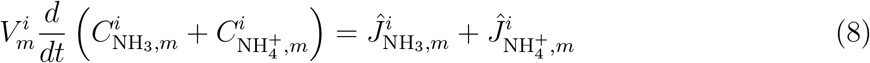

where *m* corresponds to ‘C’ or ‘P’. 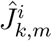 denotes the net flux of solute *k* into compartment *m*; specifically, 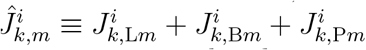.

The buffer pairs are assumed to be in equilibrium:

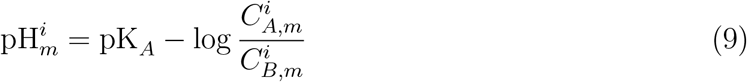

where 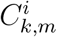 denotes the concentration of solute *k* in compartment *m*. The buffer pairs (A,B) are 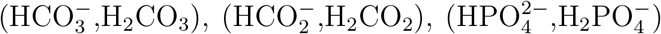, and 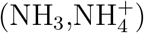. The pH of compartment *m* is given by conservation of hydrogen ion:

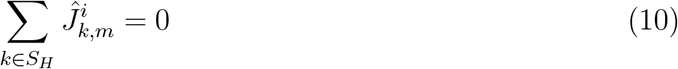

where 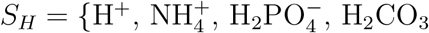, and H_2_CO_2_*}*.

## 3 Conservation equations in the lumen

Conservation of luminal fluid and non-reacting solutes is given by

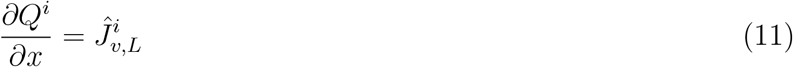

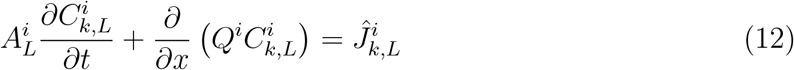

where *Q*^*i*^ denotes volume flow (per tubule) and 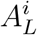 is the luminal cross-sectional area. 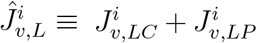 denotes overall (transcellular and paracellular) water flux and 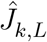 denotes the analogous solute flux.

For the reacting solutes, conservation is applied to the total buffers, assumed to be in equilibrium (Eq. 9):

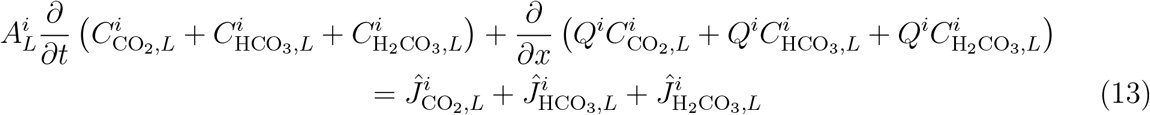

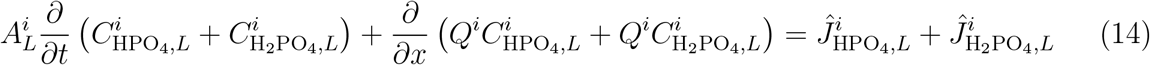

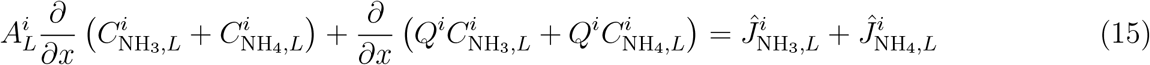

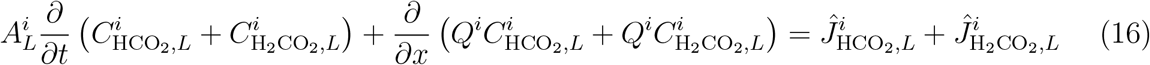

## 4 Flux calculations

Volume fluxes are calculated using the Kedem-Katchalsky equation [6]:

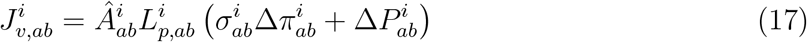

where 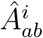 denotes a scaling factor that determine the area available for transport between compartments *a* and 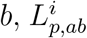 is the hydraulic permeability, 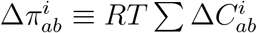 is the osmotic pressure gradient, *RT* is the product of the gas constant and thermodynamic temperature, 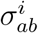 is the reflective coefficient, and 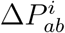 denotes the hydrostatic pressure gradient.

Transmembrane solute fluxes may include passive and active components. An uncharged solute may be driven across a membrane passively by an chemical gradient:

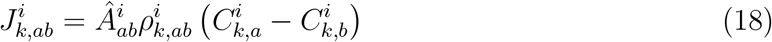

where 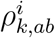 denotes membrane permeability to solute *k*.

For a charged particle, that flux may be driven by an electrochemical gradient across ion channels, described using the Goldman-Hodgkin-Katz current equation:

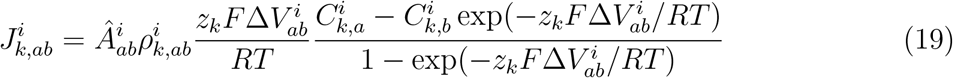

where 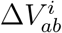 is the electrical potential gradient, *z*_*k*_ is the valance, and *F* is the Faraday’s constant. In case of an ion channel that exhibits circadian rhythms in its expression (e.g., ENaC or ROMK), the permeability of that channel 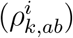 is taken to vary in time.

Solute fluxes may include additional components, such as coupled transport across cotransporters and/or exchangers and primary active transport across ATP-driven pumps [7].

## 5 Results

By applying appropriate parameters that describe kidney size, hemodynamics, and transporter expression pattern, the model can simulate the kidneys of a rat (e.g., [8, 11]), human (e.g., [9, 2]), or a mouse (e.g., [15]); male and female animals (e.g., [3, 4, 5, 14]); under special dietary conditions (e.g., [8, 1]); in an animal or patient with diabetes (e.g., [11, 13, 16]) and under SGLT2 inhibition (e.g., [11, 10]).

Given a set of model parameters, model equations can be solved to steady state to predict spredict luminal fluid flow, hydrostatic pressure, membrane potential, luminal and cytosolic solute concentrations, and transcellular and paracellular fluxes through transporters and channels. As an illustration, model predictions of a male rat kidney are shown in Fig. 2 [12].

**Figure 2:**
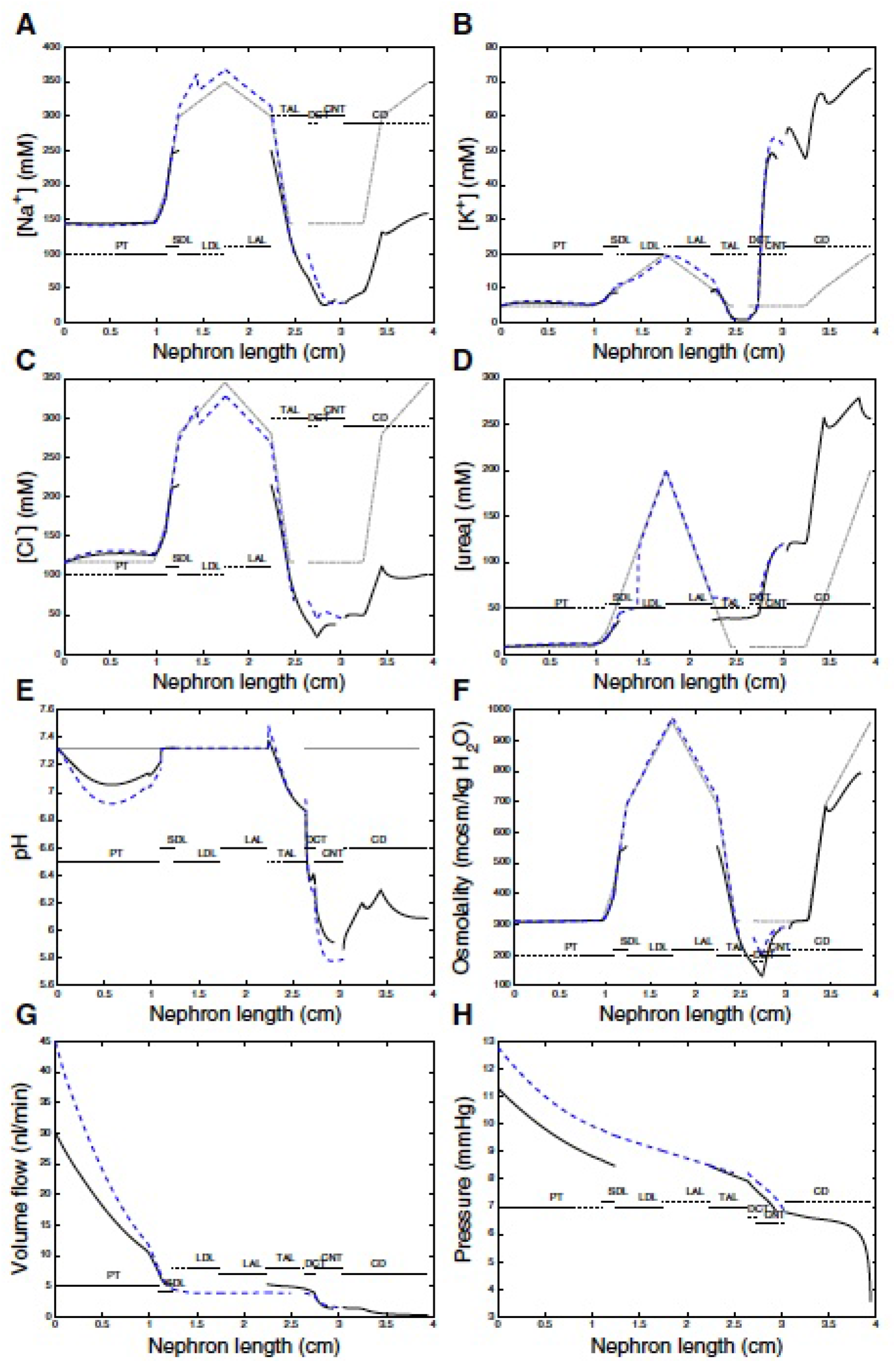
Sample model results taken from Ref. [12]. Profiles of tubular fluid solute concentrations (A–D), pH (E), osmolality (F), volume flow (G), and fluid pressure (H). Solid black lines, superficial nephron. Blue dashed line, longest juxtamedullary nephron. Grey dashed-dotted lines, interstitial fluid solute concentrations, pH, and osmolality. PT, proximal tubule; SDL, short or outer-medullary descending limb; LDL/LAL, thin descending/ascending limb; TAL, thick ascending limb; DCT, distal convoluted tubule; CNT, connecting duct; CD, collecting duct.

